# The association of ubiquitin-associated protein 2-like and Ras-GTP-activating protein SH3 domain binding protein 1 mediated by small nucleolar RNA is essential for stress granule formation

**DOI:** 10.1101/2022.04.20.488692

**Authors:** Eri Asano-Inami, Akira Yokoi, Mai Sugiyama, Toshinori Hyodo, Tomonari Hamaguchi, Hiroaki Kajiyama

## Abstract

Stress granules (SGs) are dynamic, non-membranous structures composed of non-translating mRNAs and various proteins and play critical roles in cell survival under stressed conditions. Extensive proteomics analyses have been performed to identify proteins in SGs; however, the molecular functions of these components in SG formation remain unclear. In this report, we show that ubiquitin-associated protein 2-like (UBAP2L) is a novel component of SGs. UBAP2L localized to SGs in response to various stresses, and its depletion significantly suppressed SG organization. Proteomics and RNA sequencing analyses found that UBAP2L formed a protein-RNA complex with Ras-GTP-activating protein SH3 domain binding protein 1 (G3BP1) and small nucleolar RNAs (snoRNAs). In vitro binding analysis demonstrated that snoRNAs were required for UBAP2L association with G3BP1. In addition, decreased expression of snoRNAs reduced the interaction between UBAP2L and G3BP1 and suppressed SG formation. Our results reveal a critical role of a novel SG component, the UBAP2L/snoRNA/G3BP1 protein-RNA complex, and provide new insights into the regulation of SG assembly.

## Introduction

Messenger ribonucleoprotein (mRNP) granules are non-membranous dynamic structures that contain non-translating messenger RNAs (mRNA) and proteins that regulate various stages of the life cycle of cellular mRNAs, including mRNA translation, pre-mRNA processing, localization and degradation [1]. The major types of mRNP granules are stress granules (SGs) [1,2], P-bodies (PBs) [1,3], germ granules [4] and neuronal transport granules [5]. Among these, SGs and PBs are the most well-characterized mRNP granules in cells. SGs are absent under normal growth conditions but are induced by stress, such as heat shock, osmotic shock, or oxidative stress. PBs are usually observed under normal growth conditions and are involved in RNA decay [1]. Both structures share many of the same proteins and RNAs, and both are dynamically fused and separated under stressed conditions [6]. SGs have been reported to be associated with the pathogenesis of cancer, neurodegeneration, inflammatory disorders and viral infections [7]; however, the details of SG assembly/disassembly and the resulting effects on cell signaling and survival programs remain unknown.

The stresses that induce SG organization activate one or two of four protein kinases, HRI (heme-regulated initiation factor 2 α kinase), PERK (PKR like endoplasmic reticulum kinase), PKR (protein kinase R), and GCN2 (general control nonderepressible 2). These kinases phosphorylate serine 51 of eIF2α, which inhibits formation of the eIF2/tRNAi^Met^/GTP ternary complex that is essential for translation initiation [8]. In the absence of this ternary complex, translating ribosomes are unable to function, leading to the accumulation of translationally stalled mRNPs. Some mRNP components are phosphorylated, ubiquitinated [9,10], arginine methylated [11-13], or O-GlcNAc modified [14] under stressed conditions, promoting mRNPs aggregations for SG organization. Although eIF2α phosphorylation is a major trigger for SG assembly, some chemicals have been reported to induce SG formation independent of phosphorylation, through the disruption of eIF4E interaction with eIF4G [15]. SGs are composed of polyadenylated mRNAs, translation initiation factors (eukaryotic initiation factors; eIFs), the 40S ribosomal subunit, and various RNA binding proteins (RBPs) that control mRNA structure and function [1,2]. One of the best studied RBPs in SGs is G3BP1 (Ras-GTP-activating protein SH3 domain binding protein 1), which is expressed in various tissues and conserved in a wide range of species. G3BP1 has a conserved acidic domain, a nuclear transport factor 2-like domain, several SH3 domain binding motifs, an RNA recognition motif and an RGG/RG motif that is rich in arginine and glycine [16]. G3BP1 interacts with many SG components and possesses DNA/RNA helicase activity in vitro [17]. Ser 149 of G3BP1 is dephosphorylated under stressed conditions, which is critical for self-aggregation and SG nucleation [18,19]. G3BP2, a homolog of G3BP1, is also expressed in many tissues and cell lines, and depletion of both proteins inhibited SG formation [20]. SGs also contain other RBPs, such as TDP-43, FUS, TIA1 and ATXN2, whose loss of function may lead to neurological and neuromuscular disorders [21-24]. In addition to RBPs, SGs recruit multiple enzymes, including protein kinases, phosphatases, and methyltransferases, to alter signaling pathways during the cellular adaptation to stress [9-14].

A recent mass spectrometry analysis of purified SGs revealed that SGs contains various RBPs [25]. We recently identified that ubiquitin-associated protein 2-like (UBAP2L) is a novel substrate of protein arginine methyltransferase 1 (PRMT1) and is necessary to ensure proper kinetochore-microtubule attachment for the progression of mitosis [26]. Other reports have shown that UBAP2L is a BMI-binding protein necessary for cell survival [27] and is associated with cancer progression [28,29]; however, the exact function of the protein in cells remains uncertain. UBAP2L has an RGG/RG motif that is often found in RBPs [30]. In this report, we show that UBAP2L is localized to SGs and is required for SG assembly. In addition, UBAP2L forms a complex with small nucleolar RNAs (snoRNAs) and G3BP1.

## Results

### UBAP2L localizes to SGs

To determine whether UBAP2L localizes to SGs, HeLa cells treated with 0.5 mM arsenite or 0.3 M sorbitol for 30 min were immunostained for UBAP2L together with SG marker proteins, such as G3BP1, TIAR, eIF4E and PABPC1. As shown in Figure 1a, UBAP2L accumulated into multiple small dot-like structures and colocalized with each of these marker proteins 30 min after arsenite or sorbitol treatment. Other stresses that induce SG formation, such as heat and hydrogen peroxide treatment, also promoted colocalization of UBAP2L with G3BP1 (Supplementary Fig. 1a). Some SG-localized proteins are known to aggregate in cells when exogenously overexpressed. Consistent with this, GFP-tagged UBAP2L (GFP-UBAP2L) aggregated around the nucleus with endogenous G3BP1 or eIF4E under non-stressed conditions. Similar to endogenous UBAP2L, GFP-UBAP2L accumulated in dot-like structures after 30 min of arsenite treatment (Fig. 1b and Supplementary Fig. 1b). G3BP1 overexpression has been reported to induce eIF2α phosphorylation [31], which inhibits translation initiation to promote SG organization. However, GFP-UBAP2L expression did not affect eIF2α phosphorylation (Supplementary Fig. 1c). Cycloheximide (CHX) blocks SG formation by inhibiting translation elongation [32]. To further confirm UBAP2L localization to SGs, cells were treated with arsenite in the presence or absence of CHX and immunostained for UBAP2L and G3BP1. In the presence of CHX, UBAP2L and G3BP1 localized diffusely throughout the cell and did not show any accumulation in SGs (Fig. 1c). These results clearly show that UBAP2L is a novel component of SGs. Some SG components are known to localize to PBs as well. Cells were immunostained for UBAP2L together with DCP1A, PB marker protein [1]. UBAP2L did not accumulate at the DCP1A-positive dots, indicating that UBAP2L is a specific component of SGs (Fig. 1d).

**Figure 1.**
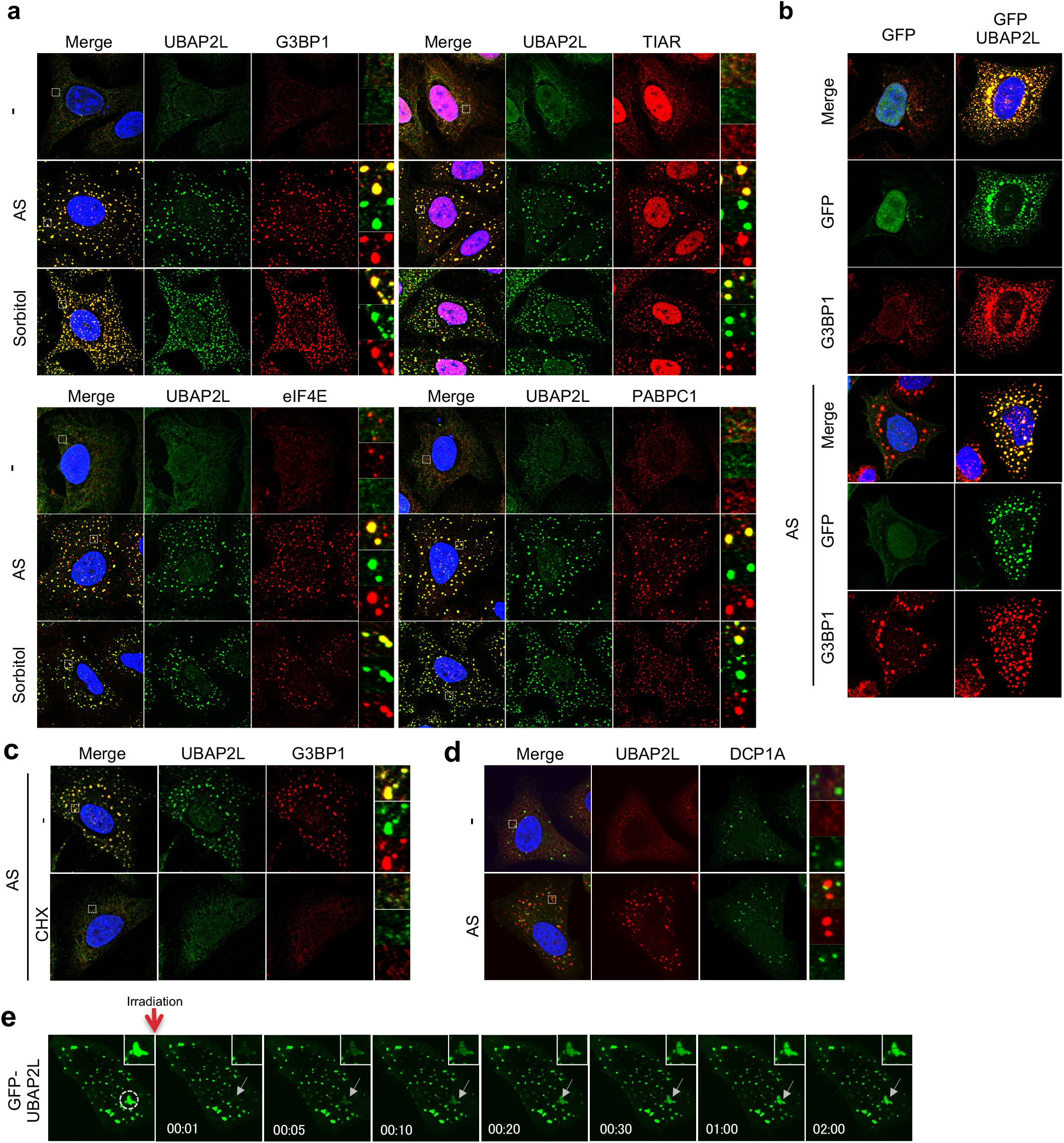
UBAP2L is localized to SGs. **a** HeLa cells treated with 0.5 mM arsenite (AS) for 30 min or 0.3 M sorbitol for 30 min were immunostained for UBAP2L together with G3BP1, TIAR, eIF4E, or PABPC1. **b** HeLa cells were transfected with plasmids encoding GFP or GFP-UBAP2L, and 24 h later, the cells were treated with or without 0.5 mM arsenite (AS) for 30 min and immunostained for GFP and G3BP1. **c** HeLa cells were treated with 0.5 mM arsenite for 30 min in the presence or absence of cycloheximide (CHX) and immunostained for G3BP1 and UBAP2L. **d** Cells treated with 0.5 mM arsenite were immunostained for DCP1A and UBAP2L. **e** Fluorescence photo bleaching recovery of a GFP-UBAP2L-expressing SG was monitored by confocal microscopy in the presence of 0.5 mM arsenite. The arrow shows the monitored SG. The time after irradiation is shown in each photograph.

SGs are dynamic structures, and many proteins associate with SGs for only a short time; these proteins rapidly shuttle between SGs and the cytoplasm [6]. We performed a FRAP assay to examine UBAP2L localization to SGs. HeLa cells that constitutively expressed GFP-UBAP2L were treated with arsenite for 30 min, and an SG was photobleached using the 405 nm laser of an FV1000 microscopy (Olympus). The diminished fluorescence of GFP-UBAP2L in the SG was recovered within 2 min (Fig. 1e), which indicates that, similar to other SG components, UBAP2L rapidly move between SGs and the cytoplasm.

### UBAP2L is essential for SG assembly

We next tested whether UBAP2L was required for SGs organization. HeLa cells transfected with two different siRNAs were treated with arsenite and immunostained for UBAP2L and G3BP1. As shown in Figure 2a, transfection of UBAP2L siRNAs clearly disrupted SG organization. Both siRNAs significantly reduced UBAP2L abundance, but the expression of other SG marker proteins, including G3BP1, TIAR, eIF4E, and PABPC1, was not affected (Fig. 2b). The disruption of SG formation can result from inhibition of stress-induced eIF2α phosphorylation. To rule out this possibility, we examined eIF2α phosphorylation and translation inhibition after arsenite treatment in the absence of endogenous UBAP2L. UBAP2L depletion did not reduce arsenite-induced eIF2α phosphorylation (Fig. 2b). Consistent with this, translation was suppressed by arsenite in UBAP2L-siRNA-transfected cells at a level similar to that of control-siRNA-transfected cells (Fig. 2c). To further confirm the critical role of UBAP2L in SG organization, UBAP2L-depleted cells were immunostained for other SG marker proteins (Fig. 2d), and the number of SGs per cell was counted. As shown in Figure 2e, f, g and h, the number of SGs in UBAP2L-depleted cells was approximately 10-20% of that in control-siRNA-transfected cells. These results show that UBAP2L is essential for SG formation, independent of eIF2α phosphorylation and translation inhibition.

**Figure 2.**
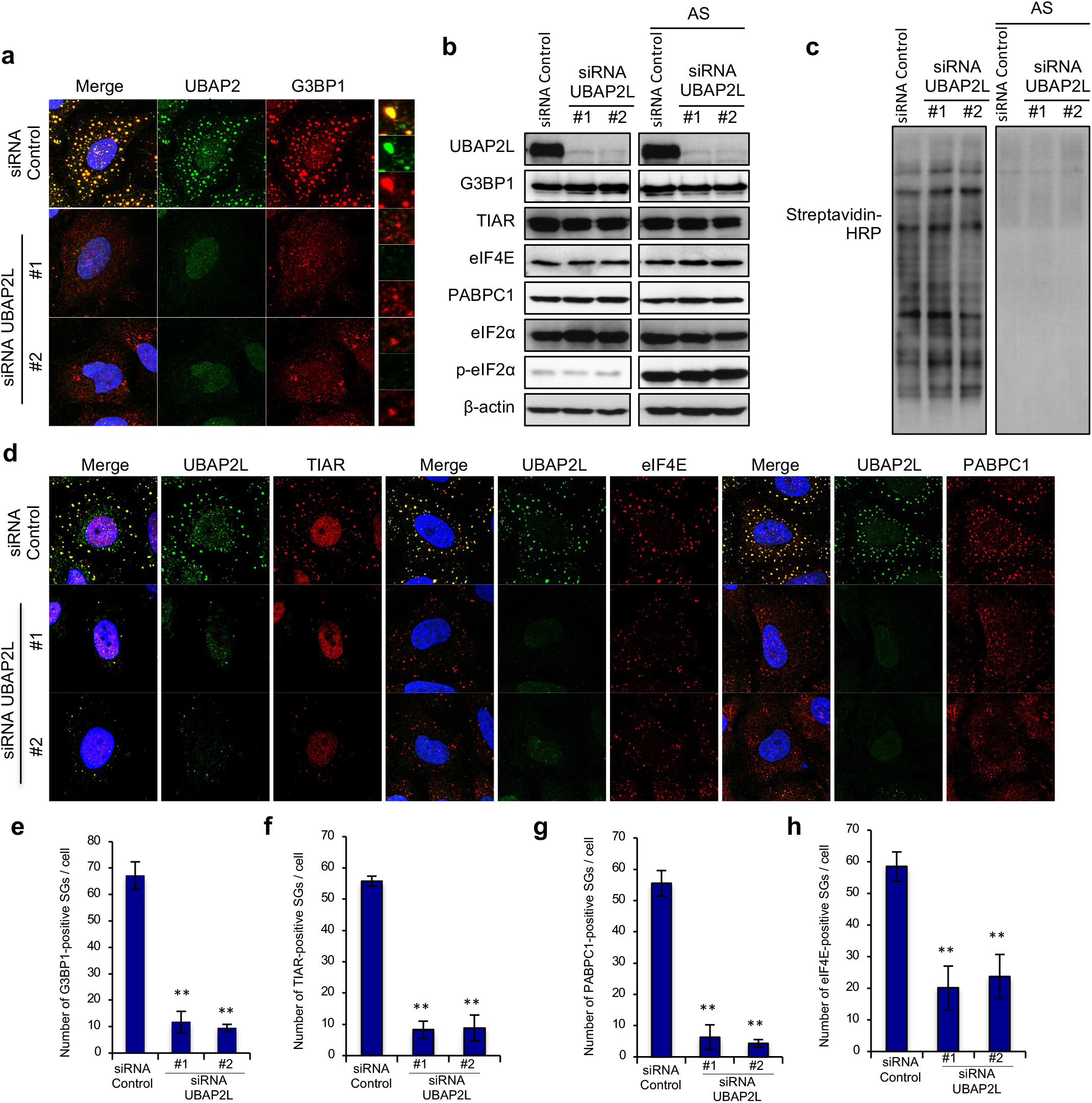
UBAP2L is essential for SG assembly. **a** HeLa cells were transfected with control or UBAP2L siRNAs, and 72 h later,the cells were treated with 0.5 mM arsenite for 30 min and immunostained with anti-UBAP2L and anti-G3BP1 antibodies. **b** Cells were transfected with the indicated siRNAs, and 72 h later, cells were treated with or without 0.5 mM arsenite for 30 min and lysed. The expression of UBAP2L, G3BP1, TIAR, eIF4E, or PABPC1 was examined by immunoblot. **c** Cells were treated with L-azidohomoalanine (AHA) with or without arsenite for 30 min, and lysed. The lysates were reacted with a Click-it Biotin Protein Analysis Detection Kit according to manufacturer’s protocols. The reacted samples were immunoblotted with an anti-streptavidin-HRP antibody. **d** Cells transfected with siRNAs were treated as in **a** and immunostained for TIAR, eIF4E, or PABPC1 and UBAP2L. **e-h** The numbers of SGs positive for each indicated SG marker per cell are presented in the graph. Three independent experiments were performed and 20 cells were evaluated for each experiment. (** *P* < 0.01).

### Amino acids 194-983 of UBAP2L are required for binding to G3BP1 and SG organization

We performed mass spectrometry analysis to search for proteins that associate with UBAP2L. In this analysis, G3BP1/2, Caprin1 and the fragile X mental retardation family proteins; FXR1/2 and FMRP, which are SG-localized proteins [18-20,33], were found to coprecipitate with UBAP2L (Fig. 3a). To confirm the association of UBAP2L with these proteins, GFP-tagged G3BP1/2, FXR1/2, FMRP or Caprin1 was transiently expressed in 293T cells together with FLAG-UBAP2L or FLAG tag. The cell lysates were immunoprecipitated with an anti-FLAG antibody, and the immunoprecipitates were subjected to immunoblot analysis. GFP-G3BP1/2, FXR1/2, FMRP and Caprin1 coecipitated with FLAG-UBAP2L but not with the FLAG tag (Fig. 3b and Supplementary Fig. 3a). Endogenous UBAP2L also coprecipitated with these proteins (Fig. 3c and Supplementary Fig. 3b), indicating that UBAP2L associates with multiple SG-localized proteins.

**Figure 3.**
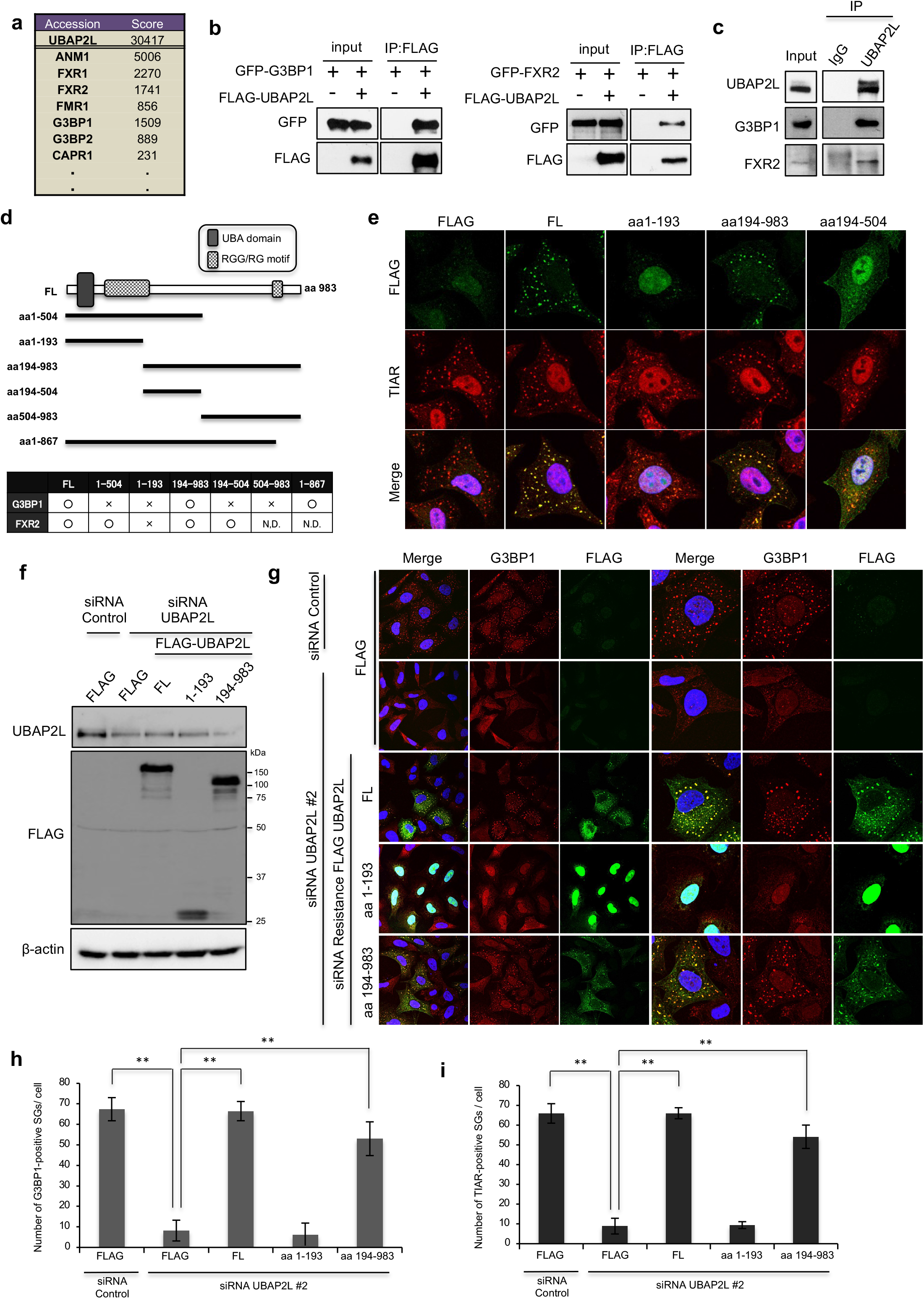
The interaction between UBAP2L and G3BP1 is required for SG assembly. **a** List of proteins identified by mass spectrometry analysis. **b** 293T cells were transfected with FLAG-UBAP2L together with GFP-G3BP1 or GFP-FXR2. After 24 h, the cells were lysed and immunoprecipitated with an anti-FLAG antibody. The immunoprecipitates were immunoblotted for GFP and FLAG. **c** HeLa cells were lysed and immunoprecipitated with either anti-rabbit IgG or anti-UBAP2L antibodies. The immunoprecipitates were blotted with anti-UBAP2L, anti-G3BP1 or anti-FXR2 antibodies. **d** Schematic representation of UBAP2L deletion constructs (N.D. (not detected)). **e** HeLa cells that constitutively expressed each construct were treated with 0.5 mM arsenite for 30 min and immunostained for TIAR and FLAG. **f** HeLa cells that constitutively expressed each siRNA resistant cDNA were depleted of endogenous UBAP2L by UBAP2L siRNA #2. The cells were lysed and immunoblotted with anti-FLAG and anti-UBAP2L antibodies. **g** Cells that constitutively expressed each constructs were depleted of UBAP2L by siRNA and immunostained for FLAG and G3BP1. **h** The number of G3BP1-positive SGs per cell is presented in the graphs (n = 20, ** *P* < 0.01). **i** The number of TIAR-positive SGs per cell is presented in the graphs (n = 20, ** *P* < 0.01).

UBAP2L has a ubiquitin-associated domain (aa50-80) and RGG/RG motifs (aa134-189, aa868-869). To determine which regions are required for the association with SG-localized proteins, FLAG-tagged UBAP2L deletion constructs were transiently expressed in 293T cells together with GFP-tagged G3BP1 or FXR2, and their association was examined by immunoprecipitation (Supplementary Fig. 3c and d). We found that the aa194-983 fragment of UBAP2L, which does not contain any specific domain, was required for the interaction with G3BP1. By contrast, the aa194-504 fragment was sufficient for the interaction with FXR2 (Fig. 3d). To assess whether these UBAP2L deletion mutants localized to SGs, HeLa cells constitutively expressing each construct were treated with arsenite and immunostained for the FLAG tag and TIAR. As shown in Figure 3e, the aa194-983 fragment, which interacts with G3BP1, localized to SGs, whereas the aa1-193 fragment, which interacts with neither G3BP1 nor FXR2, did not. A region that is sufficient for the interaction with FXR2 (aa194-504) did not show any accumulation in SGs.

We examined whether the G3BP1-binding region of UBAP2L was essential for SG formation. We generated siRNA-resistant, FLAG-tagged full-length UBAP2L (FLAG-FL-UBAP2L(R)) and aa194-983 UBAP2L (FLAG-194-984(R)) by introducing silent mutations in a region targeted by UBAP2L siRNA #2. HeLa cells that constitutively expressed the FLAG tag alone, FLAG-FL-UBPA2L(R), FLAG-194-984(R), and FLAG-1-193 (FLAG-tagged aa1-193 UBAP2L) were transfected with UBAP2L siRNA #2 (Fig. 3f) and immunostained for FLAG together with G3BP1 or TIAR (Fig. 3g). Although FLAG-1-193 did not rescue SG organization, both FLAG-FL-UBAP2L(R) and FLAG-194-983(R) significantly restored the number of SGs produced per cell (Fig. 3h and i). These results reveal that the G3BP1 interacting region of UBAP2L is essential for SG assembly.

### Small RNAs regulate the interaction between UBAP2L and G3BP1

Posttranslational modifications, such as phosphorylation and methylation, play a critical role in protein interactions. We previously reported that the N-terminal RGG/RG motif of UBAP2L was arginine methylated by PRMT1 [26]; thus, we next examined whether arginine methylation was required for the interaction between UBAP2L and G3BP1. HeLa cells were treated with or without Adox, an arginine methylation inhibitor, and the interaction was examined by immunoprecipitation. As shown in Supplementary Figure 4a, addition of Adox did not affect the interaction. In addition, UBAP2L clearly accumulated in SGs in the presence of Adox (Supplementary Fig. 4b). Mass spectrometry analysis showed that some serine/threonine residues of UBAP2L were phosphorylated. Cell lysates were immunoprecipitated with an anti-UBAP2L antibody and treated with or without lambda phosphatase (Supplementary Fig. 4c). The phosphosphatase treatment did not affect the interaction between UBAP2L and G3BP1, indicating that these modifications are not essential for the association of UBAP2L with G3BP1.

It has been reported that UBAP2L is an RBP [34]. G3BP is also known as an RBP [1-3], so we speculated that the interaction between UBAP2L and G3BP1 is regulated by RNA. HeLa cells lysates were immunoprecipitated with an anti-UBAP2L antibody and treated with Benzonase, which degrades all forms of DNA and RNA. Interestingly, the association between UBAP2L and G3BP1 was diminished by Benzonzse treatment (Fig. 4a), but the association between UBAP2L and FXR2 was not affected (Fig. 4a). We also used DNase I or RNase A to specifically degrade either DNA or RNA. Addition of DNase I did not affect the interaction (Fig. 4b) but RNase A treatment significantly reduced the interaction (Fig. 4c). These results imply that the association is dependent on the existence of RNA.

**Figure 4.**
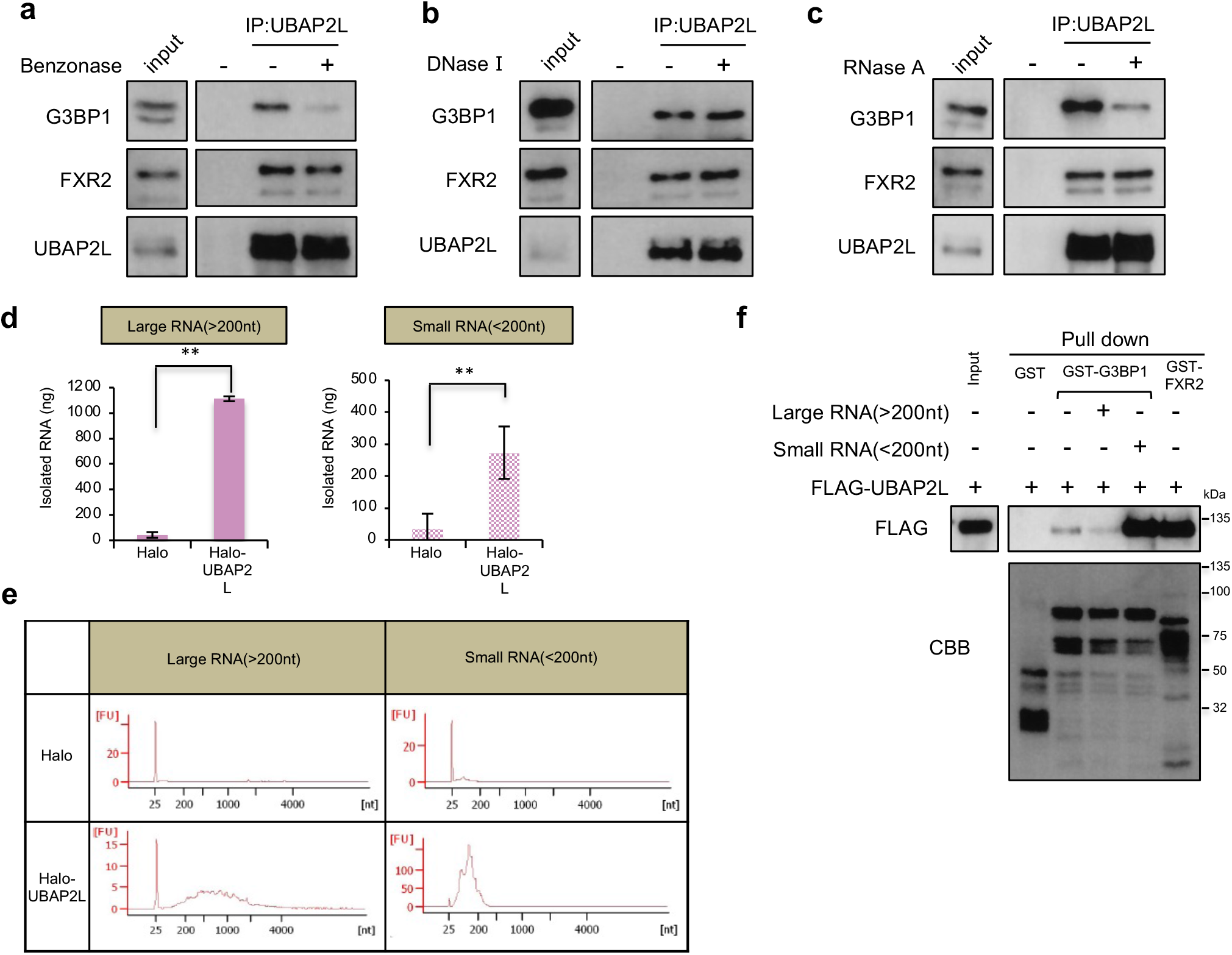
Small RNAs are required for the interaction between UBAP2L and G3BP1. **a** UBAP2L was immunoprecipitated with an anti-UBAP2L antibody from HeLa cell lysates. The immunoprecipitates were treated with Benzonase and immunoblotted with anti-UBAP2L, anti-G3BP1 and anti-FXR2 antibodies. **b.c** HeLa cells were lysed and immunoprecipitated as in **a**. The immunoprecipitates were treated with DNase 1 or RNase A and immunoblotted with the indicated antibodies. **d** 293T cells that constitutively expressed the Halo tag or Halo-UBAP2L were immunoprecipitated with an anti-Halo antibody. Large RNAs (>200 nt) or small RNAs (<200 nt) were isolated from the immunoprecipitates. The graphs show the amounts of isolated RNAs (ng). Three independent experiments were performed (** *P* < 0.01). **e** The Bioanalyzer quality analysis of each isolated RNAs is presented in the graphs. **f** Immunoprecipitaed FLAG-UBAP2L was mixed with recombinant GST, GST-G3BP1 or GST-FXR2 bound to glutathione beads with or without larges RNAs(20 μg) or small RNAs (8 μg). The beads were precipitated and subjected to immunoblot analysis with an anti-FLAG antibody. CBB indicates Coomassie brilliant blue staining of recombinant proteins.

To determine if UBAP2L binds to RNA, we performed a RIP assay. 293T cells that constitutively expressed the Halo tag or Halo-UBAP2L were immunoprecipitated, and large RNAs (>200nt) or small RNAs (<200nt) were isolated. Both large and small RNAs were specifically isolated from precipitates of Halo-UBAP2L-expressing cells (Fig. 4d). The Agilent Bioanalyzer was used to confirm the size and quality of each of isolated RNA. These results show that UBAP2L is an RBP (Fig. 4e).

We examined whether large or small RNAs were required for the interaction between UBAP2L and G3BP1. FLAG-UBAP2L was transiently expressed in 293T cells and purified by immunoprecipitation using an anti-FLAG antibody, followed by extensive washing with RIPA buffer and high-salt buffer to eliminate any binding proteins and RNAs. The purified FLAG-UBAP2L was incubated with GST, GST-G3BP1, or GST-FXR2, which were purified from bacteria, at 4°C in the presence or absence of 20 *μ*g of large RNAs or 8 *μ*g of small RNAs. GST, GST-G3BP1 and GST-FXR2 were precipitated by glutathione agarose beads, and the precipitates were subjected to immunoblot with an anti-FLAG antibody. FLAG-UBAP2L associated with GST-FXR2 in the absence of RNA. By contrast, FLAG-UBAP2L was coprecipitated only in the presence of small RNAs (Fig. 4f). These results show that small RNAs are required for the interaction between UBAP2L and G3BP1 and that RNA is dispensable for the UBAP2L -FXR2 interaction.

### UBAP2L and G3BP1 form a complex with snoRNAs

To determine which small RNAs mediate the interaction between UBAP2L and G3BP1, small RNAs isolated by RIP using Halo-UBAP2L and Halo-G3BP1 precipitates were analyzed by RNA sequencing. We obtained approximately 50 million reads, and 14% were uniquely mapped to UCSC human genome 19. The RNAs with the top 10 RPM scores are shown in Figure 5a and Supplementary Figure 5a. A circle graph shows the classification of RNAs with the top 100 RPM scores (Fig. 5b and Supplementary Fig. 5b). Interestingly, 60-70% of the top 100 RNAs in each result was small nucleolar RNA (snoRNA). SnoRNA is approximately 60-300 nucleotides long and is necessary for modification of precursor ribosomale RNA (rRNA) [35]; however, there has been no report indicating a role for snoRNAs in SG organization. SnoRNAs are classified into two groups, C/D box and H/ACA box snoRNAs. These two groups of RNAs have distinctive sequences, but some regions are evolutionally conserved [36]. Most of the snoRNAs that we obtained by RNA sequencing analysis were C/D box snoRNAs (SNORDs) (Fig. 5a and b, Supplementary Fig. 5a and b). To confirm the association between snoRNAs and UBAP2L, the Halo tag, Halo-UBAP2L, Halo-G3BP1 or Halo-CPEB was expressed in 293T cells and precipitated using Halo beads. RNA was isolated from each precipitate and subjected to qRT-PCR analysis to detect SNORD44 and SNORD49A, which are C/D box snoRNAs. CPEB1 is an SG-localized protein that does not interact with UBAP2L. The analysis showed that both SNORD44 and SNORD49A bound to UBAP2L and G3BP1, but not CPEB1 (Fig. 5c). Furthermore, we carried out an in vitro binding assay using purified proteins and in vitro-transcribed SNORD44. Full length SNORD44 was cloned into the pBluescript vector, and both the sense and anti-sense SNORD44 sequences were transcribed in vitro. In the presences SNORD44 sence sequence, FLAG-UBAP2L clearly associated with GST-G3BP1 (Fig. 5d). By contrast, the anti-sense sequence of SNORD44 did not promote the interaction (Fig. 5d). We also examined other SNORDs, including SNORD49A, SNORD30 and SNORD56. The RPM score of SNORD56 was significantly lower than those of SNORD30 and SNORD49A. As demonstrated in Figure 5e, UBAP2L interacted with G3BP1 in the presence of these snoRNAs; however, the SNORD56-mediated interaction was less significant than the SNORD30- or SNORD49A-mediated interaction. These results indicate that some specific C/D box snoRNAs form a complex with UBAP2L and G3BP1.

**Figure 5.**
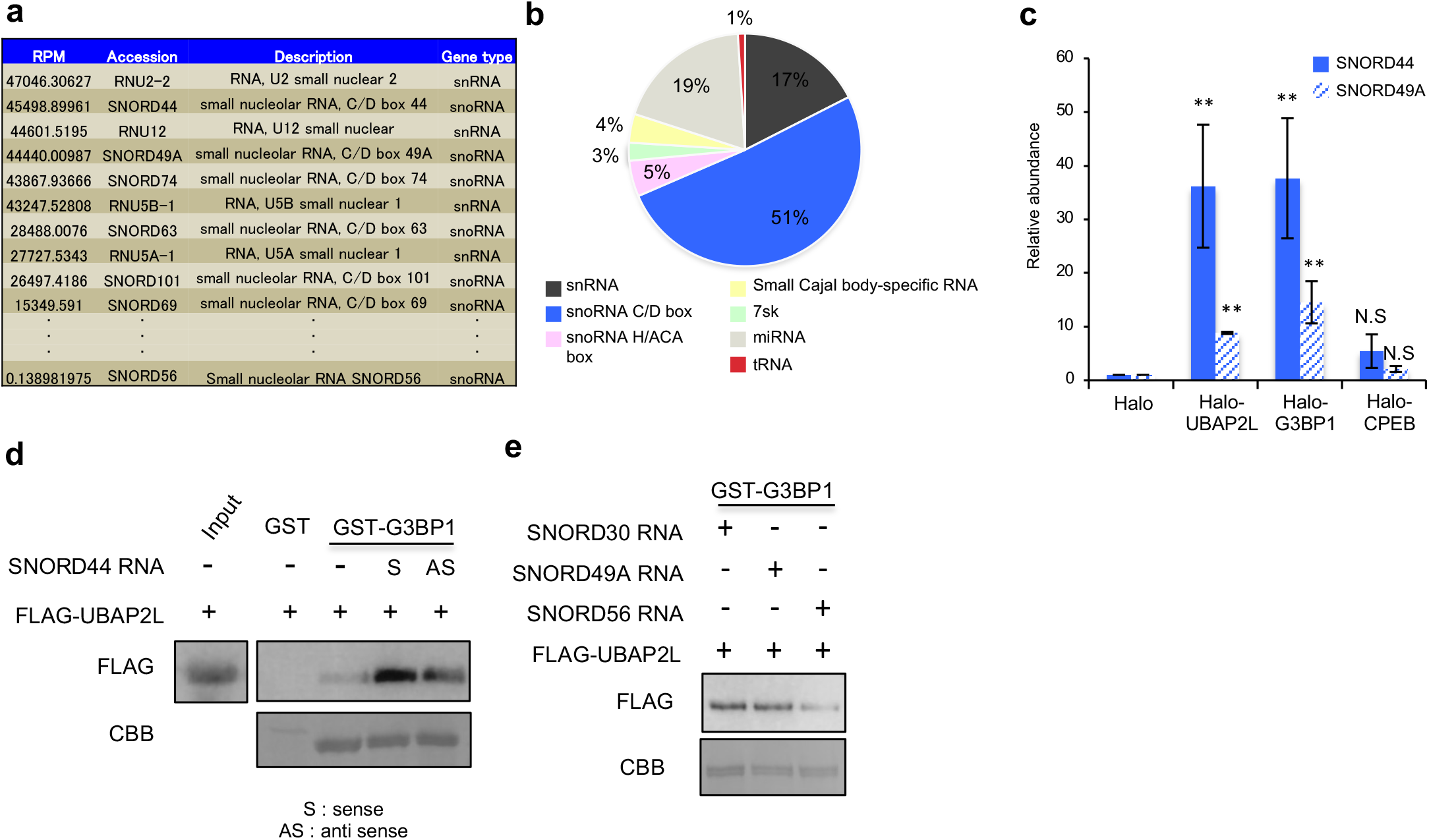
C/D box snoRNAs mediates the association between UBAP2L and G3BP1. **a** The small RNAs with the top 10 UBAP2L RPM scores are presented (n = 2). **b** The percentages of each type of small RNAs with the top 100 UBAP2L RPM scores are shown in the cicle graph (n = 2). **c** Small RNAs were isolated from immunoprecipitates from the lysate of 293T cells that constitutively expressed the Halo tag, Halo-UBAP2L, Halo-G3BP1 or Halo-CPEB. The levels of SNORD44 and SNORD49A were examined by quantitative RT-PCR (qRT-PCR) (n = 3, ** *P* < 0.01, N.S. (not significant) *P* > 0.05). **d** Immunoprecipitated FLAG-UBAP2L was mixed with recombinant GST or GST-G3BP1 and incubated with or without in vitro-transcribed SNORD44 sense (S) or anti-sense (AS) sequences. Protein-RNA complex precipitated by glutathione agarose beads were immunoblotted with an anti-FLAG antibody. CBB indicates Coomassie brilliant blue staining of recombinant proteins. **e** In vitro-transcribed SNORD30, SNORD49A or SNORD56 were mixed with immunoprecipitated FLAG-UBAP2L and GST-G3BP1 and subjected to precipitation with glutathione agarose beads followed by immunoblot analysis. Both UBAP2L and G3BP1 RPM scores of SNORD56 is significantly lower than that of SNORD30 and SNORD49A.

### C/D box snoRNAs regulats SGs assembly

We next investigated whether C/D box snoRNAs were required for SG formation. Most C/D box snoRNAs are processed from spliced introns to become functional for rRNA modifications. NOP56, NOP58, NHP2L1 and fibrillarin are critical factors in the generation of C/D box snoRNAs [37]. To reduce the abundance of C/D box snoRNAs, HeLa cells were transfected with NOP56 or NOP58 siRNA. Transfection of siRNAs significantly decreased NOP56 and NOP58 mRNA expression (Fig. 6a). The abundance of SNORD49A was significantly suppressed by transfection of either siRNA, whereas SNORD44 expression was reduced only by NOP58-depletion (Fig. 6b).

**Figure 6.**
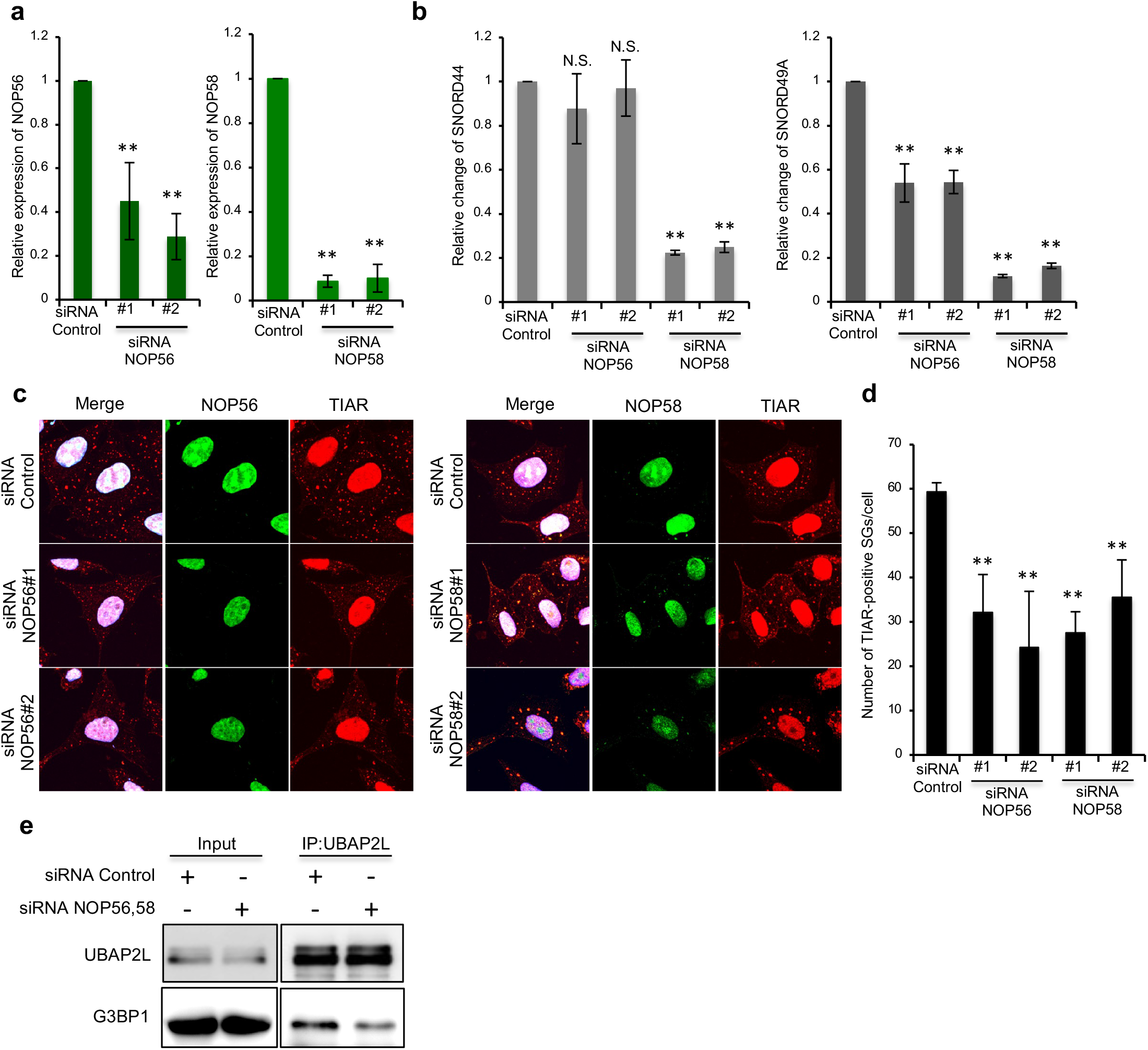
C/D box snoRNAs regulate SG assembly. **a** HeLa cells were transfected with control, NOP56 or NOP58 siRNAs. After 72 h, the expression of NOP56 (left graph) or NOP58 (right graph) mRNA was examined by qRT-PCR. Three independent experiments were performed (** *P* < 0.01). **b** The abundance of SNORD44 (left graph) or SNORD 49A (right graph) was examined by qRT-PCR (n = 3, ** *P* < 0.01, N.S. (not significant) *P* > 0.05). **c** HeLa cells were transfected with control, NOP56, or NOP58 siRNAs, and 72 h later, the cells were treated with 0.5 mM aresenite for 30 min and immuostained with anti-TIAR together with anti-NOP56 or anit-NOP58 antibodies. **d** The number of SGs per cell are presented in the graph. Three independent experiments were performed (n = 20, ** *P* < 0.01). **e** HeLa cells were transfected with control siRNA or a combination of NOP56 #1 and NOP58 #1 siRNAs, and 72 h later, the cells were lysed and immunoprecipitated with anti-UBAP2L antibody. The immunoprecipitates were subjected to immunoblot analysis.

**Figure 7.**
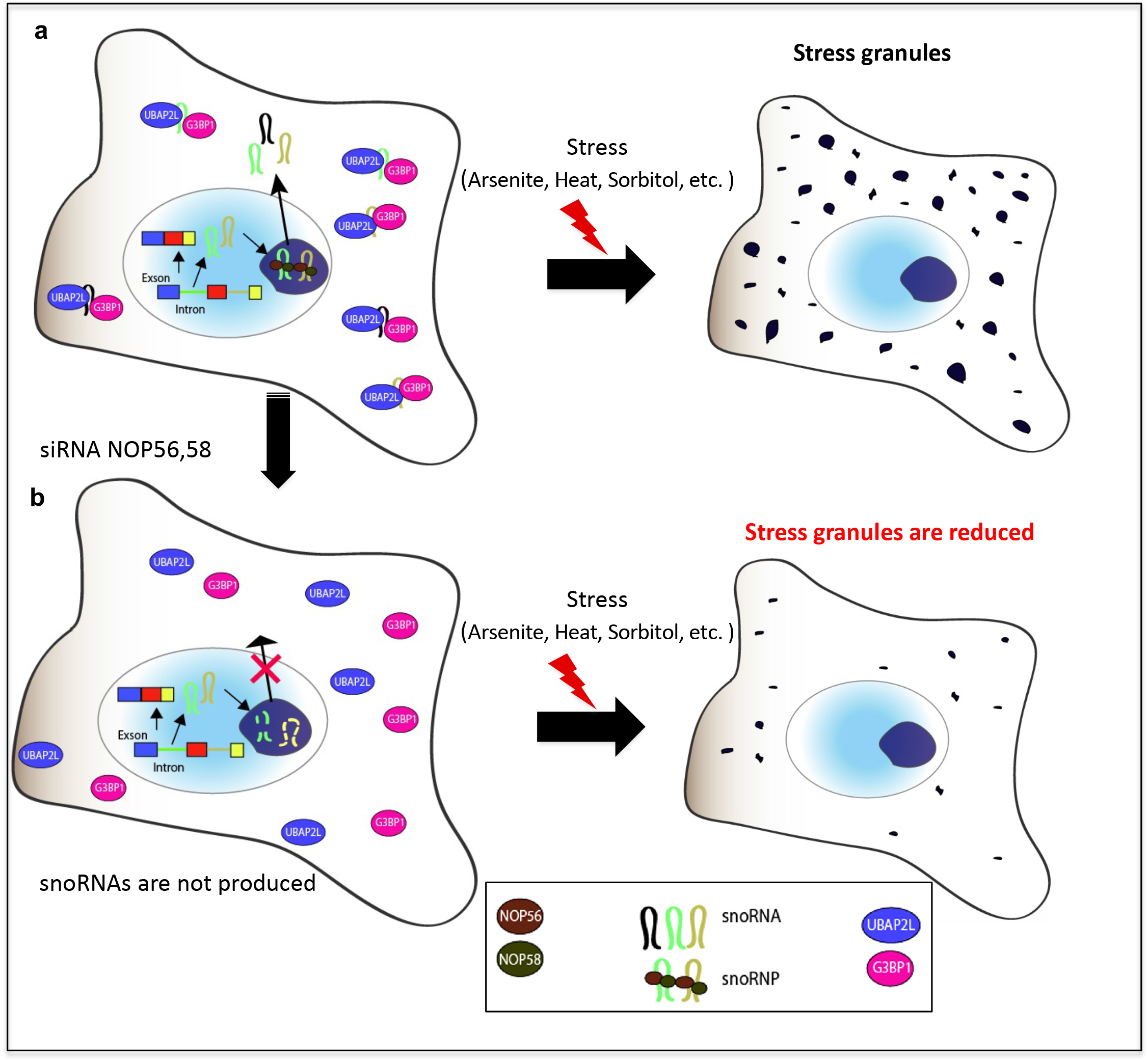
A schematic model of the UBAP2L-snoRNA complex in SG. **a** In the nucleus, snoRNAs are produced by introns and complex with NOP56 and NOP58 as snoRNPs and matured by endonucleases. Some snoRNAs are produced in the cytoplasm. UBAP2L and G3BP1 complex with C/D box snoRNAs in the cytoplasm. Once cells are stimulated by stress, SGs are formed normally. The cells can adjust to the stress conditions to survive. **b** When the expression of C/D box snoRNAs is reduced by NOP56 or NOP58 depletion, UBAP2L and G3BP1 cannot form a complex. The number of SGs is reduced, and it is difficult for the cell to adjust to stress.

We next examined whether SG assembly was suppressed by either NOP56 or NOP58 depletion. SGs were organized in cells depleted of NOP56 or NOP58 (Fig. 6c); however, the number of SGs per cell in cells transfected with either NOP56 or NOP58 siRNA was significantly lower than that in control siRNA-transfected cells (Fig. 6d). Finally, we examined the association between UBAP2L and G3BP1 in cells transfected with both NOP56 and NOP58 siRNAs. HeLa cells transfected with both NOP56 siRNA #1 and NOP58 siRNA #1 were lysed and immunoprecipitaed with an anti-UBAP2L antibody. As shown in Figure 6e, the interaction between UBAP2L and G3BP1 was suppressed in NOP56- and NOP58-depleted cells. Collectively, these results show that C/D box snoRNAs mediate the association between UBAP2L and G3BP1 and promote SGs assembly under stress conditions.

## Discussion

In this report, we identified a novel role for UBAP2L in SG organization. Endogenous as well as exogenously expressed GFP-UBAP2L clearly accumulated in SGs in response to various stresses. Knockdown of UBAP2L by two different siRNAs significantly reduced the number of SGs per cell, and exogenous expression of siRNA-resistant UBAP2L restored SG formation in cells depleted of endogenous UBAP2L. These results clearly show that UBAP2L is a novel component of SGs and essential for SG assembly. UBAP2L knockdown did not affect stress-induced eIF2α phosphorylation or translation inhibition, indicating that UBAP2L does not affect pathways leading to translational repression. In addition, expression of other SG-localized proteins was not reduced by UBAP2L depletion. Thus, it is likely that UBAP2L is required for SG assembly. Our mass spectrometry analysis further supported the importance of UBAP2L in SG organization. UBAP2L formed a complex with other SG-localized proteins, including FMR1, G3BP1, and Caprin1. UBAP2L is conserved in a wide range of species, and the Drosophila homolog is called Lingerer. A previous study reported that Lingerer interacted with FMR1, Caprin and Rasputin, a Drosophila homolog of G3BP1 [38]. Although whether Lingerer is required for SG formation remains to be investigated, it was shown that Lingerer regulates the JAK/STAT signaling pathway to restrict cell proliferation during development [39]. The functions of mammalian UBAP2L have yet to be studied, but recent studies showed that UBAP2L is associated with cancer cell proliferation [28,29]. Therefore, mammalian UBAP2L may be involved in processes other than SG formation, such as cell proliferation, by regulating signaling pathways.

Our RIP assay and in vitro binding assay revealed that UBAP2L association with G3BP1 was mediated by snoRNAs. There are two types of snoRNAs, C/D box snoRNAs and H/ACA box snoRNAs. Interestingly, most of the snoRNAs obtained by RNA sequencing were C/D box snoRNAs. Depletion of proteins that are required for the generation of mature C/D box snoRNAs significantly suppressed UBAP2L association with G3BP1 and SG formation. These results indicate that C/D box snoRNAs are critical for the assembly of the UBAP2L/snoRNA/G3BP1 protein-RNA complex to promote SG organization. C/D box snoRNAs are 60-100-nucleotide-long transcripts characterized by conserved box C (RUGAUGA, where R is a purine) and box D (CUGA) motifs [40]. The major function of C/D box snoRNAs is 2’-*O*-methylation of their target rRNAs; however, recent studies have revealed that C/D box snoRNAs have additional functions [41,42]. For example, SNORD60 is involved in intracellular cholesterol trafficking [43], and SNORDs U32a, U33 and U35a mediate lipotoxic stress, possibly through their cytosolic function [44]. Furthermore, SNORD55A and SNORD55B have been shown to bind and inhibit K-Ras and suppress tumorigenesis [45]. These reports and our study show that C/D box snoRNAs have multiple functions other than rRNA modification.

Accumulating studies have revealed that not only mRNAs but also miRNAs are involved in SG assembly [46,47]. The localization of mRNA to SGs has been shown by fluorescence in situ hybridization (FISH) using dT probes [23]; however, miRNA localization to SGs has not been confirmed. We could not detect C/D box snoRNA localization to SGs by FISH using multiple probes for SNORD44 (Supplementary Fig.4c) and SNORD49A (data not shown). One reason we could not observe snoRNA localization to SGs may be the methodological difficulties of FISH analysis of small RNA, such as insufficient sensitivity or inability of the probes to access their target snoRNAs. Another possibility is that snoRNAs are processed into shorter RNAs in SGs. It has been shown that snoRNAs are processed into shorter RNAs called sno-miRNAs, which are over 18 nt in length and functions are unknown [48-53]. SGs contain RNA endonucleases, such as Argonaute family proteins [46,47], which may shorten snoRNAs in SGs and cause them to be undetectable by FISH. Immunoprecipitation analysis showed that the UBAP2L/snoRNA/G3BP1 complex is formed in the absence of stress, thus, snoRNAs may be necessary for the initial phase of SG assembly and once SGs are formed, snoRNAs in the complex may be cleaved by RNA endonucleases for other functions. Although our study shows that snoRNAs are important for SG organization, further investigations are necessary to confirm snoRNA localization to SGs and understand the mechanisms of snoRNA-mediated SG assembly and cell survival.

In summary, we have shown that UBAP2L forms a protein-RNA complex with G3BP1 and snoRNAs and that these complex is essential for SG assembly. UBAP2L also associates with additional SG-localizing proteins, such as FMR1, FXR1, FXR2, and Caprin1. In addition, UBAP2L interacts with not only small RNAs but also mRNAs. RNA sequencing demonstrated that several mRNAs encoding apoptosis-related proteins are associated with UBAP2L; thus, future studies elucidating the physiological roles of UBAP2L will provide more insight into the molecular mechanisms and relevance of SGs and cell survival.

## Methods

### Cells, antibodies, and chemicals

HeLa and 293T cells were purchased from RIKEN BRC (Tsukuba, Japan) and cultured DMEM containing 10% FBS at 37°C. To generate an anti-UBAP2L antibody, a fragment including amino acid (aa) residues 513-660 of UBAP2L fused with glutathione S-transferase (GST) was produced in bacteria, and recombinant protein was purified by using glutathione agarose beads (Sigma-Aldrich, St. Louis, MO, USA). The protein was mixed with Freund’s adjuvant (Sigma-Aldrich) and injected into a rabbit 4 times, every 2 weeks. To purify the anti-UBAP2L antibody, we used Hi-Trap N-hydroxysuccinimide (NHS)-activated HP columns (GE Healthcare Bio-Sciences), coupled with recombinant GST-UBAP2L (aa 513-660). Other antibodies were obtained from the following companies: anti-G3BP1 (611126), anti-TIAR (610352) and anti-FXR2 (61130) antibodies were obtained from BD Biosciences (San Jose, CA, USA); anti-G3BP1 (A301-033A) and anti-G3BP2 (A302-040A) antibodies were obtained from Bethyl Laboratories (Montgomery, TX); anti-eIF2α (5324), anti-phospho-eIF2α(3398), anti-FXR1 (12295),anti-FMRP (4317), and anti-streptavidin-HRP (3999) antibodies were obtained from Cell Signaling (Danvers, CA, USA); anti-eIF4E (sc-9976) and anti-PABPC1 (sc-32318) antibodies were obtained from Santa Cruz Biotechnology (Santa Cruz, CA, USA); anti-DCP1A antibody (H00055802-M06), Abnova (Taipei, Taiwan); anti-FLAG antibody (014-22383), Wako (Osaka, Japan); anti-GFP (598) and anti-DIG (M227-3) antibodies were obtained from MBL (Nagoya, Japan); anti-Halo antibody (G921A) were obtained from Promega (Madison, WI, USA); and then anti-ADMA antibody were obtained from Active Motif (Carlsbad, CA, USA).

### siRNA transfection

The following control siRNA and siRNAs used to suppress UBAP2L, G3BP1/2, FXR2, and NOP56/58 expression were purchased from Thermo Fisher Scientific (Waltham, MA USA): *Silencer*^®^ Select siRNA Control (4390843); and *Silencer*^®^ Select Pre-designed siRNA UBAP2L #1(s19176) and #2(s230223), G3BP1 #1 (s19754) and #2 (s19755), G3BP2 #1 (s19206) and #2 (s19207), FXR2 #1 (s18243) and #2 (s18244), NOP56 #1 (s20642) and #2 (s20643), and NOP58 #1 (s28390) and #2 (s28391). HeLa cells were transfected with 5 _n_M siRNA using Lipofectamine RNAiMAX (Invitrogen, Carlsbad, CA. USA).

### Immunofluorescence analysis

Cells cultured on fibronectin-coated glass cover slips were transfected with siRNAs or vectors and then stimulated with 0.5 mM arsenite for 30 min, 300 mM sorbitol for 30 min, 1 mM hydrogen peroxide for 1.5 h, or heat (44°C) for 30 min. Cells were fixed with 4% paraformaldehyde for 10 min on ice and permeabilized with PBS containing 0.5% Triton for 5 min. The cells were blocked with PBS containing 7% FBS for 30 min and incubated with primary antibodies for 1 h. After washing with PBS, the cells were incubated with Alexa Fluor 488- or Alexa Fluor 594-labeled secondary antibodies (Invitrogen) and Hoechst (Dojindo) for 1 h. Images were acquired using an FV1000 laser scanning confocal microscope (Olympus, Tokyo, Japan).

### Fluorescence recovery after photobleaching (FRAP)

Images were recorded using an FV1000 laser scanning confocal microscope (Olympus). HeLa cells constitutively expressing GFP-UBAP2L were cultured on a glass-bottom dish and treated with 0.5 mM arsenite for 20 min. An SG was irradiated with a laser 405 nm for 10 sec, images were scanned every second for 5 min after bleaching.

### RNA immunoprecipitation (RIP) assay

293T cells constitutively expressing the Halo tag, Halo-UBAP2L or Halo-G3BP1 were lysed in lysis buffer (25 mM Tris-HCl (pH 7.6), 150 mM NaCl, and 0.1% NP-40) supplemented with a phosphatase inhibitor cocktail (Nacalai Tesque, Inc., Japan), protease inhibitor (Promega), 1.5mM DTT, and RNase inhibitor (Takara) for 10 min on ice and treated with DNaseI (Takara) for 10 min at room temperature. The lysates were centrifuged at 15,000 rpm for 10 min and the supernatants were incubated with Halo Resin (Promega) for 12 h. The resin beads were washed with wash buffer (25 mM Tris-HCl (pH 7.6), 150 mM NaCl, and 0.005% NP-40) five times, and suspended in ML buffer (NucleoSpin miRNA: Takara) to isolate large and small RNAs.

### Small RNA-sequencing

Small RNAs isolated by RIP were assessed with a Nanodrop 2000 instrument (Thermo Fisher Scientific) and RNA integrity was determined with an Agilent 2100 Bioanalyzer (Agilent, Japan). Small RNA libraries were generated using NEB Next Small RNA Library Prep Set for Illumina (NEB). Sequencing was performed on HiSeq 1500 instrument (Illumina). The sequencing reads were trimmed using FASTX-Tool kit 0.0.13 and mapped to human reference genome hg 19 (UCSC) and the RPM (reads per million mapped reads) was calculated using Strand NGS ver 2.1 (Agilent Technologies).

### Statistical analysis

Three independent experiments were performed, and the results were compared using Student’s t-test. The data are represented as the means ± standard deviation (s.d).

## Acknowledgement

We are grateful to Dr. Takeshi Senga for helpful discussions and comments on the manuscript. This study was financially supported through JSPS KAKENHI Grant Numbers 21H03075, Princess Takamatsu Cancer Research Fund, Daiichi Sankyo Foundation of Life Science, the Uehara Memorial Foundation. Moreover, this study was also supported by Program for Promoting the Enhancement of Research Universities as young researcher units for the advancement of new and undeveloped fields at Nagoya University.

## Notes

### Competing Interest Statement

The authors have declared no competing interest.

