## Supplementary figure legends for "The association of ubiquitin-associated protein 2-like and Ras-GTP-activating protein SH3 domain binding protein 1 mediated by small nucleolar RNA is essential for stress granule formation"

Supplementary Figure 1

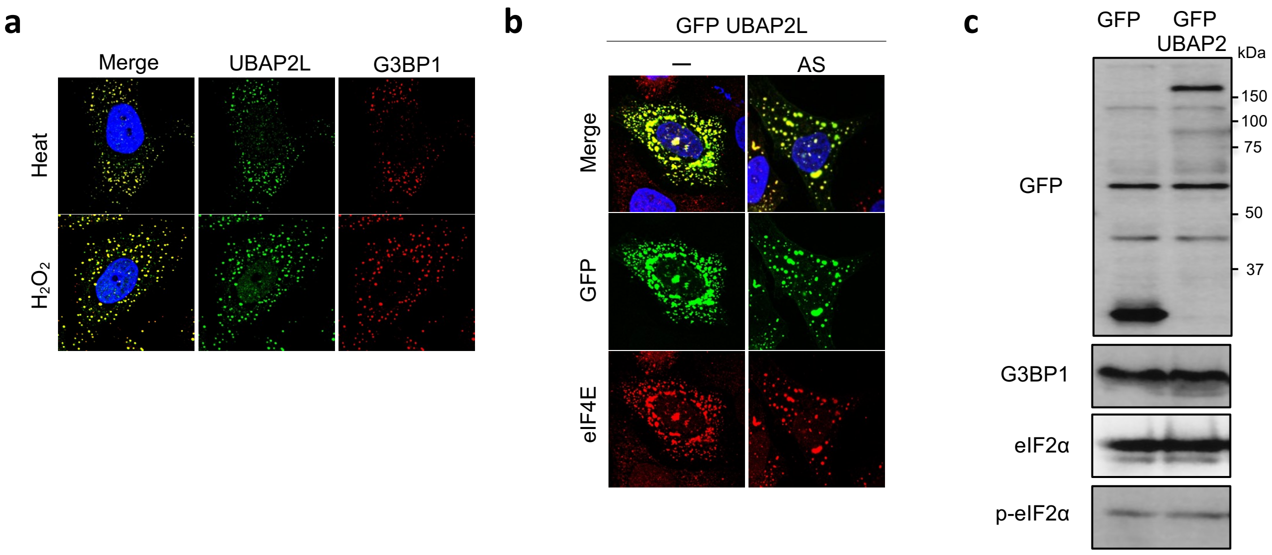

Supplementary Figure 2

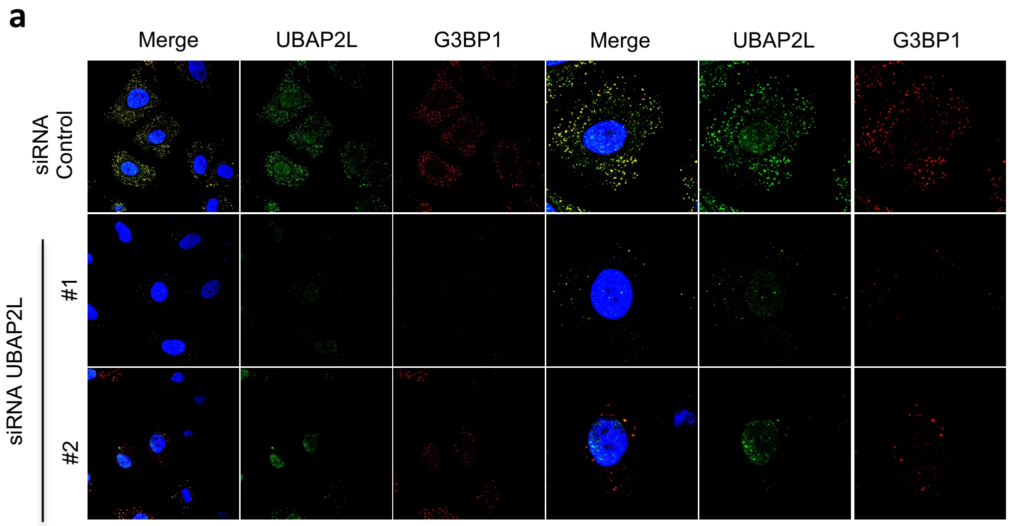


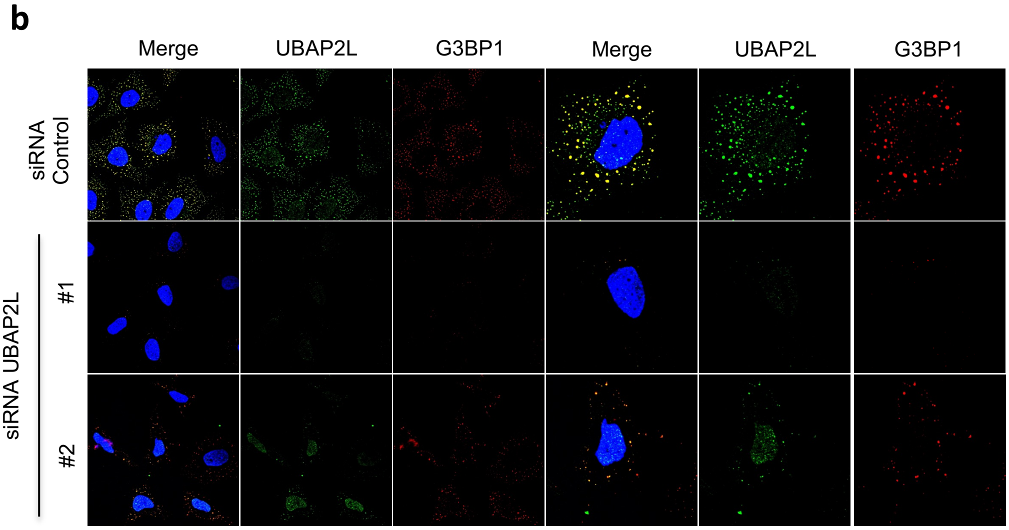

Supplementary Figure 3

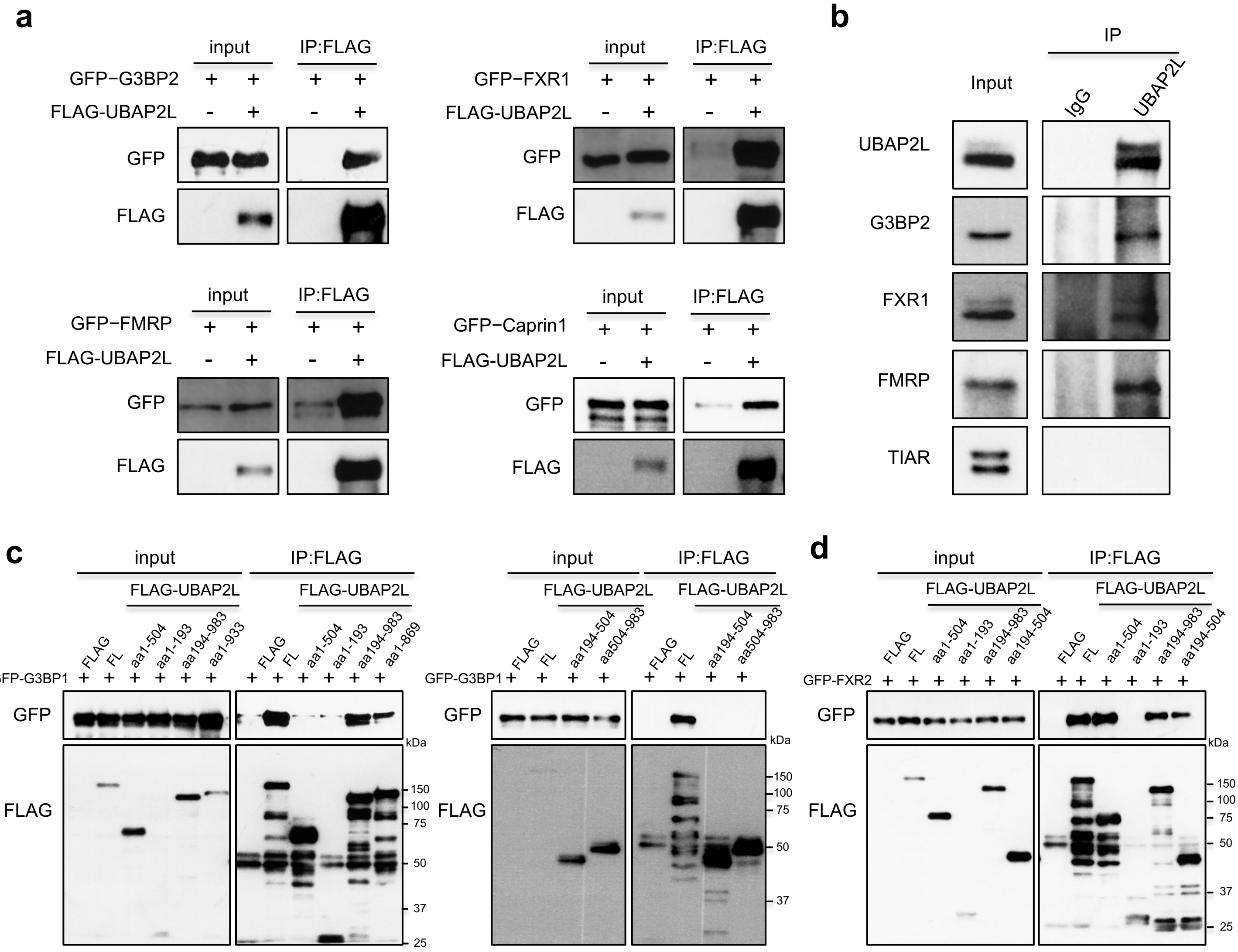

Supplementary Figure 4

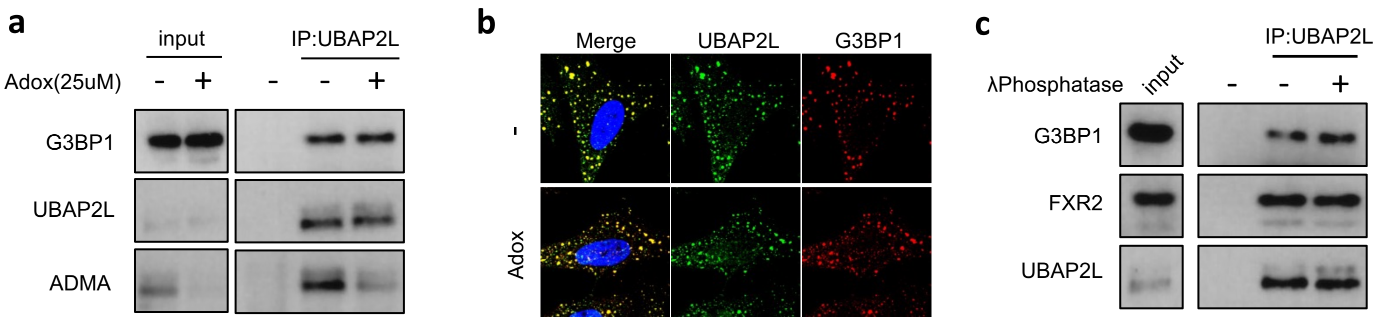

Supplementary Figure 5

**a**

| RPM(G3BP1) | Accession | Description | Gene Type |
|---|---|---|---|
| 61985.91399 | RNU2−2 | RNA, U2 small nuclear 2 | snRNA |
| 60072.56429 | SNORD44 | small nucleolar RNA, C/D box 44 | snoRNA |
| 59113.22164 | SNORD49A | small nucleolar RNA, C/D box 49A | snoRNA |
| 57704.62064 | SCARNA20 | small Cajal body−specific RNA 20 | snoRNA |
| 54777.50507 | RNU12 | RNA, U12 small nuclear | snRNA |
| 50410.84196 | SNORD74 | small nucleolar RNA, C/D box 74 | snoRNA |
| 45382.56323 | SNORD3A | small nucleolar RNA, C/D box 3A | snoRNA |
| 44895.95561 | RNU5B−1 | RNA, U5B small nuclear 1 | snRNA |
| 32952.7265 | SNORD3C | small nucleolar RNA, C/D box 3C | snoRNA |
| 32092.62619 | RNU5A−1 | RNA, U5A small nuclear 1 | snRNA |
| ⋮ | ⋮ | ⋮ | ⋮ |
| 0 | SNORD56 | Small nucleolar RNA, C/D box 56 | snoRNA |

**b**

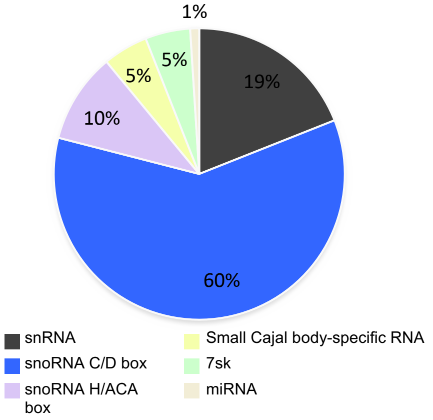


**c**

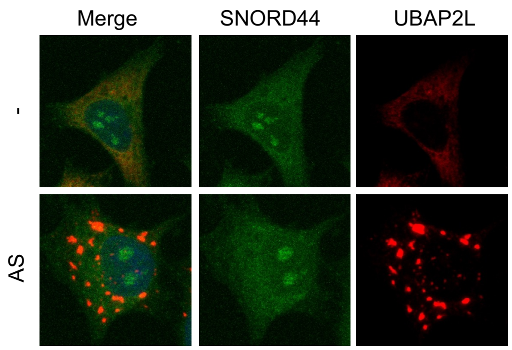
