## Supplemental 0420 for "The association of ubiquitin-associated protein 2-like and Ras-GTP-activating protein SH3 domain binding protein 1 mediated by small nucleolar RNA is essential for stress granule formation"

Supplementary Figure legends

Supplementary Figure 1. UBAP2L is localized to SGs in response to various stress conditions. Related to Figure1. **a** HeLa cells were treated with heat shocked (44℃) for 30 min or 1 mM hydorogen peroxide ($H_2O_2$) for 1.5 h. The cells were immunostained for UBAP2L and G3BP1. **b** HeLa cells were transfected with plasmids encoding GFP or GFP-UBAP2L, and 24 h later, the cells were immunostained for GFP and eIF4E with or without 0.5 mM arsenite (AS) treatment for 30 min. **c** Cells were transfected with plasmids encoding indicated cDNA and 24 h later, the cells were lysed and immunoblotted.

Supplementary Figure 2. UBAP2L depletion suppressed SG assembly by heat and sorbitol treatments. **a.b** HeLa cells were transfected with control or UBAP2L siRNAs. After 72h, the cells were treated with heat shocked (44℃) for 30min (**a**) or 0.3 M sorbitol for 30min (**b**) and immunostained with anti-UBAP2L and anti-G3BP1 antibodies.

Supplementary Figure 3. UBAP2L interacted with multiple SG localizing proteins. **a** 293T cells were transfected with FLAG-UBAP2L together GFP-G3BP2, GFP-FXR1, GFP-FMRP or GFP-Caprin1, and 24 h later, the cells were lysed and immunoprecipitated with an anti-FLAG antibody. The immunoprecipitaes were immunoblotted by anti-GFP and anti-FLAG antibodies. **b** HeLa cells were lysed and immunoprecipitaed with control or anti-UBAP2L antibodies. The immunoprecipitates were subjected to immunoblot analysis. **c** 293T cells were transfected with GFP-G3BP1 together with FLAG-UBAP2L deletion mutants. After 24 h, cells were lysed and immunoprecipitated with an anti-FLAG antibody and subjected to immunoblotting to probe for GFP or FLAG. **d** 293T cells were transfected with GFP-FXR2 together with GFP-UBAP2L deletion mutants. After 24 h, cells were lysed and immunoprecipitated with an anti-FLAG antibody and subjected to immunoblotting to probe for GFP and FLAG.

Supplementary Figure 4. Adox and phosphatase treatments did not affect the interaction between UBAP2L and G3BP1. **a** HeLa cells were cultured in the presence or absence of 25uM of Adox for 24 h, and lysed, and immunoprecipitated with an anti-UBAP2L antibody. The immuneipitates were immunoblotted with indicated antibodies. **b** Cells were cultured with or without Adox for 24 h, and then treated with 0.5 mM arsenite for 30 min. The cells were immunostained for G3BP1 and FXR2. **c** HeLa cells were immunoprecipitated with an anti-UBAP2L antibody, the immunoprecpitates were treated with lambda phosphatase and immunoblotted.

Supplementary Figure 5. The results of small RNA sequencing of G3BP1 RIP. **a** The RNAs with the top 10 G3BP1 RPM scores are presented. **b** The percentages of each type of gene of the RNAs with top 100 G3BP1 RPM scores are shown in the circle graph. **c** HeLa cells were treated with or without 0.5 mM arsenite for 30 min. **c** Cells were fixed and hybridized SNORD 44–DIG probe for overnight, and then cells were immunostained for DIG and UBAP2L.
